# Two Independent and Highly Efficient Open Source TKF91 Implementations

**DOI:** 10.1101/033191

**Authors:** Nikolai Baudis, Pierre Barbera, Sebastian Graf, Sarah Lutteropp, Daniel Opitz, Tomáš Flouri, Alexandros Stamatakis

## Abstract

In the context of a master level programming practical at the computer science department of the Karlsruhe Institute of Technology, we developed and make available two independent and highly optimized open-source implementations for the pair-wise statistical alignment model, also known as TKF91, that was developed by Thorne, Kishino, and Felsenstein in 1991. This paper has two parts. In the educational part, we cover teaching issues regarding the setup of the course and the practical and summarize student and teacher experiences. In the scientific part, the two student teams (Team I: Nikolai, Sebastian, Daniel; Team II: Sarah, Pierre) present their solutions for implementing efficient and numerically stable implementations of the TKF91 algorithm. The two teams worked independently on implementing the same algorithm. Hence, since the implementations yield identical results —with slight numerical deviations— we are confident that the implementations are correct. We describe the optimizations applied and make them available as open-source codes in the hope that our findings and software will be useful to the community as well as for similar programming practicals at other universities.

## 1 Introduction

In [8], Thorne, Kishino, and Felsenstein presented a method for pair-wise alignment of DNA sequences using a maximum likelihood (ML) approach. They developed an explicit statistical model of evolution that uses statistical insertions, deletions, and substitutions of nucleotides as basic operations for comparing two DNA sequences. An evolutionary model, that is given as input, determines at which rate these three evolutionary events occur. These rates are then used to design a dynamic programming (DP) algorithm that computes the ML pair-wise alignment between two DNA sequences.

As substitution model, we assume the standard F81 stochastic model of nucleotide substitution [4]. The algorithm computes the optimal maximum likelihood sequence alignment using three matrices, one for each evolutionary event (i.e., substitution, deletion, and insertion). The algorithm is given by the following dynamic programming recurrence:

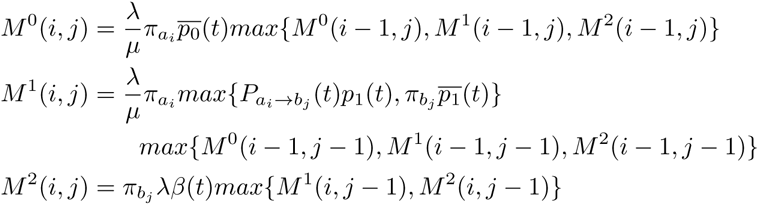

Where λ and *μ* denote the birth and death rate, π = (π*_A_*, π*_C_,* π*_G_,* π*_T_*) denotes the equilibrium frequency of the four nucleotides, *P_x→y_* the transition probability from state *x* to *y,* and 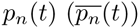the probability that after time *t* a so-called mortal link has exactly *n* descendants and one of these *is* (resp. *is not*) the original mortal link. As already mentioned, the values for *t*, *λ*, *μ*, and π are given as input. Finally, *β*(*t*) is defined as:

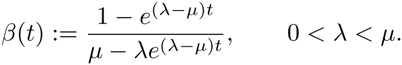

Note that, the values 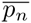can be pre-computed in constant time. For more details, please refer to the original paper [4] and the on-line task specification (see http://www.exelixis-lab.org/web/teaching/practical15/description/tkf91.pdf).

To the best of our knowledge, there is a limited amount of related work on optimizing the algorithm. In 2000 Hein *et al*. [5] presented several techniques for optimizing the algorithm. However, their focus was slightly different. One main goal of the work was to conduct a ML estimate of the model parameters. This was achieved by implementing a banded DP that only fills DP cells within a band along the diagonal of the DP matrix. Evidently, this can yield suboptimal results, but was deemed sufficient to estimate the ML parameters. Once this was done, the authors performed a full, unbanded DP computation with the previously computed ML parameter estimates. Note that, optimizing ML model parameters or using banded approaches was not part of the task specification for the students. Finally, the authors propose similar simplifications of the recursion as we do here, with the only difference that they use the term tabulation instead of lookup table or memoization we use here. Also, the authors do apparently not use a transformation into log space as we do here. Unfortunately, it was not possible to obtain a copy of the code to conduct comparisons to our implementations (pers. comm with J. Hein in August 2015).

In addition there also exists an R package that implements the TKF91 model (see http://cran.us.r-project.org/web/packages/TKF/TKF.pdf), but only for amino acid data. We did therefore not compare it to our implementations.

Finally, the Handle package [6] comprises a TKF91 implementation (see https://github.com/ihh/dart/tree/master/src/tkf) that is used for constructing multiple sequence alignments in a Bayesian setting. Since the TKF91 calculations are embedded into a MCMC (Markov Chain Monte Carlo) framework, it was not feasible to compare the performance of this TKF91 implementation with our implementations. Form a visual code inspection, it seems though, that the implementation of TKF91 is rather straight-forward, that is, not particularly optimized.

In the rest of the paper we refer to the dynamic programming paradigm as DP, and to dynamic programming matrices as DPM. The source codes developed by teams I and II are available for download under the GNU GPL license (see https://github.com/nbaudis/bioinf2015).

The remainder of this paper is organized as follows: In Section 2, we describe the teaching setup and goals. The teams then present their implementations in Sections 3 and 4, respectively. The corresponding experimental results by both teams are presented in Section 5. In the following Section 6, we summarize our teaching experiences. We conclude in Section 7.

## 2 Teaching Perspective, Goals and Course Outline

### 2.1 Teaching Setup & Goals

In the first semester of the Bioinformatics module that spans two semesters, we teach a lecture called “Introduction to Bioinformatics for Computer Scientists”, since KIT does not offer a stand-alone Bioinformatics degree. This lecture covers basic topics such as an introduction to molecular biology, classic pair-wise sequence alignment, BLAST, de novo and by-reference sequence assembly, multiple sequence alignment, phylogenetic inference, MCMC methods, and population genetics.

In the second semester of the module, students can choose if they want to do a seminar presentation or the programming practical whose results we describe here. The goal of the practical is to carry out a self-contained project and write, as well as release software, that will be useful to the evolutionary biology community. Another key focus is on using tools (e.g., static analyzers, memory checkers) that increase software quality. Note that, at a CS department, designing “classic” bioinformatics analysis pipelines using scripting languages is typically not considered as “real programming” by the students. Hence, we needed to define a project that required coding in `C/C++` or `Java`. One should also strive to avoid having the students extend existing software, since this is generally frustrating and hinders creativity.

We thus decided to ask the students to implement efficient, sequential versions of the TKF91 algorithm that can also be used as library routines. Since the DP algorithm as such, is relatively straight-forward to implement for a computer science student, the main focus was on code optimization. The students were thus asked to implement a highly efficient version of the code in `C` or `C++` using all capabilities of a modern CPU (e.g., SSE3 and AVX intrinsics). The main motivation for this, was to give the students enough time to experiment with low-level optimization strategies on modern CPUs. Moreover, this project allowed the students to apply a broad range of skills acquired in the Bioinformatics and other master-level modules at our department. Furthermore, the TKF91 model required understanding and applying the discrete pair-wise sequence alignment methods and likelihood-based models for sequence evolution introduced in the lectures. To foster competition among the teams, an award (dinner payed by A.S.) was announced for the team that would implement the fastest code. To allow for a fair comparison of the codes and the CPU-specific optimizations we provided the students access to a reference machine with 4 physical cores (Intel i7–2600 running at 3.40GHz) and 16GB RAM.

In terms of project documentation, students are usually required to write a report. However, in the present case, we jointly took the decision to write a paper about the practical and upload it to `biorxiv`. This has the positive effect that students also learn how to write scientific papers.

### 2.2 Code Quality Assessment

In order to continuously monitor and improve the quality of our source codes, we used several tools and methods throughout our project. We deployed both, static, as well as dynamic code analysis tools.

**Static Analyses** Static analyses help to identify programming errors such as incorrect programming language syntax or typing errors. We compiled our codes with the `gcc` compiler using *all* available and reasonable warning flags^3^. This allowed us to identify potential programming errors at an early stage. Due to its more pedantic nature (i.e., ability to detect more errors), we also used the `clang` compiler with respective flags^4^ periodically alongside of `gcc` to further reduce the amount of potential programming errors. Note that, `clang` conducts a static code analysis.

**Dynamic Analyses** These analyses cover runtime issues, mainly memory leaks. We used `valgrind` and its sub-module `memcheck` for detecting memory-related errors in our programs.

## 3 Implementation of Team I

Before describing our vectorized implementation, we will cover the necessary prerequisites. In Section 3.1 we describe the DP algorithm in more detail as well as the corresponding wave-front parallelism we exploited in our implementation. In Section 3.2, we describe our memory layout concept, which is required to improve the efficiency of the vectorization. Thereafter, in Section 3.3, we describe how we cache intermediate computations. Then, we explain our vectorization approach in Section 3.4. Section 3.5 outlines our considerations regarding the memory alignment of the data that constitutes an important part of the vectorized implementation.

### 3.1 Dynamic Programming and Wave-front Parallelism

DP is a widely-used technique to efficiently solve a class of, at first sight, apparently hard computational problems by breaking them down into a collection of simpler subproblems. The solutions to these subproblems are combined to reach an overall solution. Pair-wise sequence alignment falls into this class. An advantage of using DP to compute pair-wise sequence alignment is that it is comparatively straight-forward to parallelize computations along the anti-diagonals of the DPM. This DPM parallelization approach is known as wave-front parallelism. The underlying idea is that matrix entries along the same DP anti-diagonal *d* can be computed independently from each other (and hence in parallel) if the preceding anti-diagonal *d* − 1 has been computed. Therefore, we can deploy vector intrinsics to accelerate calculations along anti-diagonals.

### 3.2 Memory Layout

We first introduce our memory layout for the three DP matrices (*M*^0^, *M*^1^, *M*^2^) since it is performance-critical. Our vectorization needs to attain high data locality to efficiently use the CPU cache. To this end, we permuted the indexing scheme of the DP matrix such that neighboring entries on an anti-diagonal are stored contiguously in memory. Figure 1 depicts the memory mapping of the matrix. It is layed out neither in a column-major nor row-major fashion, but stored linearly by anti-diagonals.

**Fig. 1.**
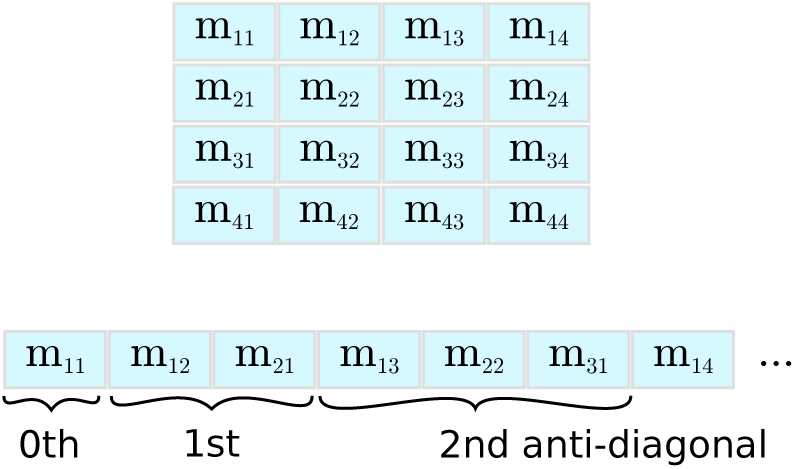
DPM memory layout

We index the matrices using a modified version of the *Cantor pairing function,* where a tuple is assigned to an offset. The original Cantor pairing function *π* assigns an integer to a pair of integers (e.g., *π* : (ℕ, ℕ) → ℕ). Since the size of the DPM is given by the length of the input sequences and because it is not infinite, we modified the pairing function to accept a tuple of two integers from the interval [0.length(Sequence 1|2)] as input. As a consequence, the DPM is divided into three parts: the *opening, intermediate,* and *closing part.* In the opening part, the modified pairing function is identical to the original function; in the intermediate and closing parts, the modified pairing function is calculated by subtracting an offset from the original pairing function. We calculate the offset via the original Cantor pairing function.

In Figure 2, we show the index calculations for the anti-diagonals of a 5 × 3-matrix. The blue frame denotes the actual boundaries of this 5 × 3 DP matrix, the blue cells represent the opening part, the green cells inside the frame the intermediate part, and the red cells inside the frame the closing part. Green and red cells outside the matrix boundaries represent the offset calculation for the intermediate and closing part. For the DP calculations, only indices that correspond to DPM entries are required. Note that, the index function has constant time and space complexity. This is essential for reducing the overhead when accessing elements in the DPMs.

**Fig. 2.**
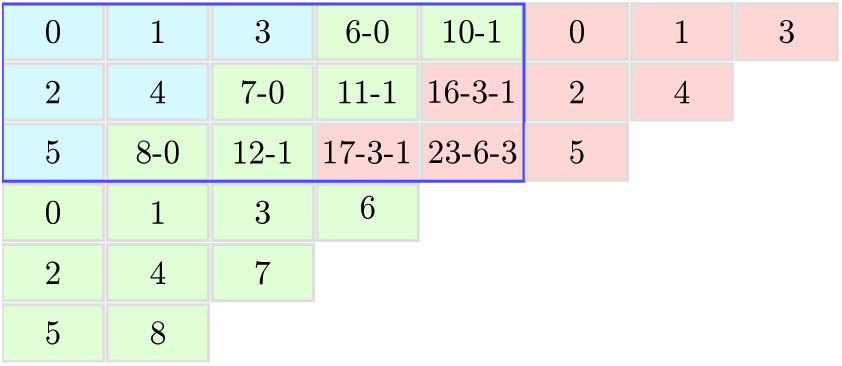
Indexing scheme for anti-diagonals

Given the anti-diagonal indexing scheme for a single matrix, we can now devise the memory layout for the three DPMs (*M*^0^, *M*^1^*, M*^2^). This is because calculating a DP value requires accessing values from all three matrices, which are located in different memory regions. To improve data locality while, at the same time, using efficient vector load and store operations, the data has to be arranged accordingly. Since we are using double-precision floating-point numbers for all calculations, an SSE3 vector (128 bit) can hold 2 values, while an AVX vector (256 bit) can hold 4 values. To this end, we evaluated the following three alternative matrix layouts (see Figure 3), where `n` always is the number of elements in one single DP matrix and `m = n / v` is the number of vectors per matrix for a vector width of `v`.

*Struct of Arrays* (*SoA*) (`struct { double m0[n], m1[n], m2[n]; } data;`) This layout allows for simple vector load and store operations. However, the three DPMs are located in separate memory regions, which decreases data locality and hence cache efficiency.

**Fig. 3.**
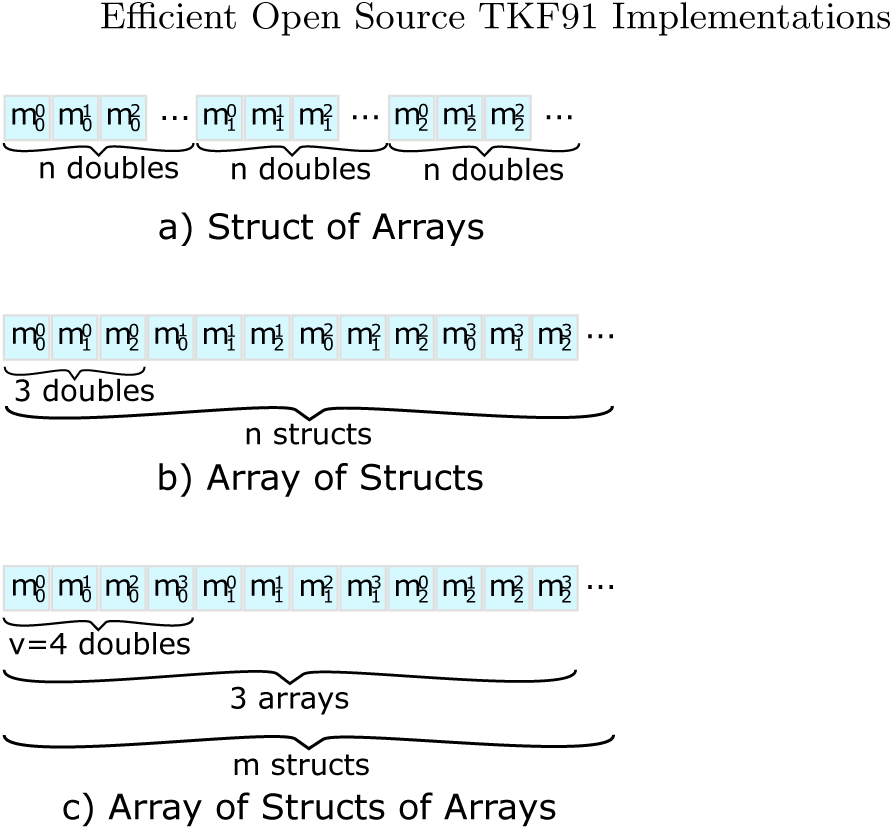
Alternative data layouts for storing anti-diagonals

*Array of Structs* (*AoS*) (`struct { double m0, m1, m2; } data[n];`) This layout exhibits improved data locality by grouping values closely together that are – in most cases – accessed simultaneously. On the other hand, loading and storing vectors requires disentangling and interleaving them, which requires several, potentially costly, vector shuffle operations.

Figure 4 illustrates these shuffle operations. For a particular position on an anti-diagonal, we use 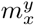to denote that element *m* of a DP matrix with index *x* is stored in slot *y* of the SSE3/AVX vector. The first row of the Figure depicts the *AoS* data layout. Initially, we perform a vector load operation to load the data into temporary vector registers. Thereafter, we shuffle the temporary vectors such that (i) each vector only contains elements from one of the three DP matrices and (ii) the individual elements are in the correct order with respect to the anti-diagonal. For both, SSE3 as well as AVX instructions, we need to perform three load operations and three `shuffle/permute` operations. For AVX, three additional blend operations are required to correctly order the data elements.

**Fig. 4.**
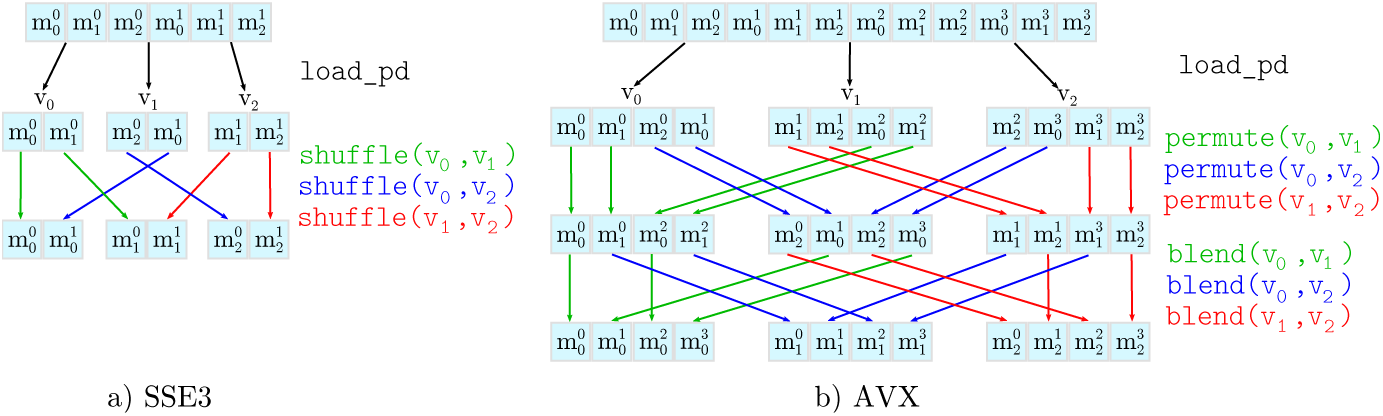
Disentangling operations for a) SSE3 (128 bit) and b) AVX (256 bit) vectors.

*Array of Structs of Arrays* (*AoSoA*) (`struct { double m0[v], m1[v], m2[v]; } data[m];`) This layout stores vectors such that adjacent elements for each of the three matrices can be directly loaded into an SSE3/AVX vector. While solving both aforementioned problems (low data locality and costly vector operations), this complicates DP cell accesses above the current anti-diagonal. The required values above the current anti-diagonal are not stored contiguously and therefore we either need to use `shuffle/permute` operations or conduct partial loads (i.e., load single elements into vectors).

All three layouts exhibit different performance characteristics with respect to cache efficiency and complexity of the load, store, as well as shuffle operations. We implemented prototypes for all three approaches to analyze their performance and found that the AoS approach performs best. Note that, the computational cost of the `shuffle` operations outweighed the improved cache efficiency in the AoS and AoSoA layouts (also see Table 5.1 in Section 5.2).

### 3.3 Memoization

The TKF91 [8] model has several parameters. Thus, it initially seemed inevitable to carry out the non-trivial computations for filling DP cells from scratch for each individual cell. However, after analyzing which expressions are constant and can thus be pre-computed and reused (i.e., memoized), we reduced the operations required for calculating a DP cell value to just three: indirection, summation, maximum.

Obvious savings can be achieved for expressions that only depend on the time parameter *t*, the birth rate λ, and the death rate *μ*. These values are constant and given as input parameters. We also observed that the cell updates in the DP algorithm only require a constant number of common sub-calculations/components: When the nucleotide states at the current indices for the two sequences are available, the score (cell value) can be computed by only using the current nucleotide pair and the neighboring values in the three DP matrices.

The memoization scheme (lookup table) for all possible configurations is shown in Figure 5. The penalty functions *C^i^* take one or two nucleotide states as parameters. Thus, these penalties can be stored in a memoization table of 4 and 16 entries, respectively. Using only 24 memoized values, we were able to substantially simplify and accelerate the cell updates. In addition, this simplification now allows to apply the logarithm for preventing numerical underflow. This was not possible before, since taking the logarithm of the original equation would have been too expensive computationally. In addition, vectorizing the cell updates is simpler, since the remaining calculations are less complex. Finally, this simplification reduces the number of possible memory layout and vectorization options.

**Fig. 5.**
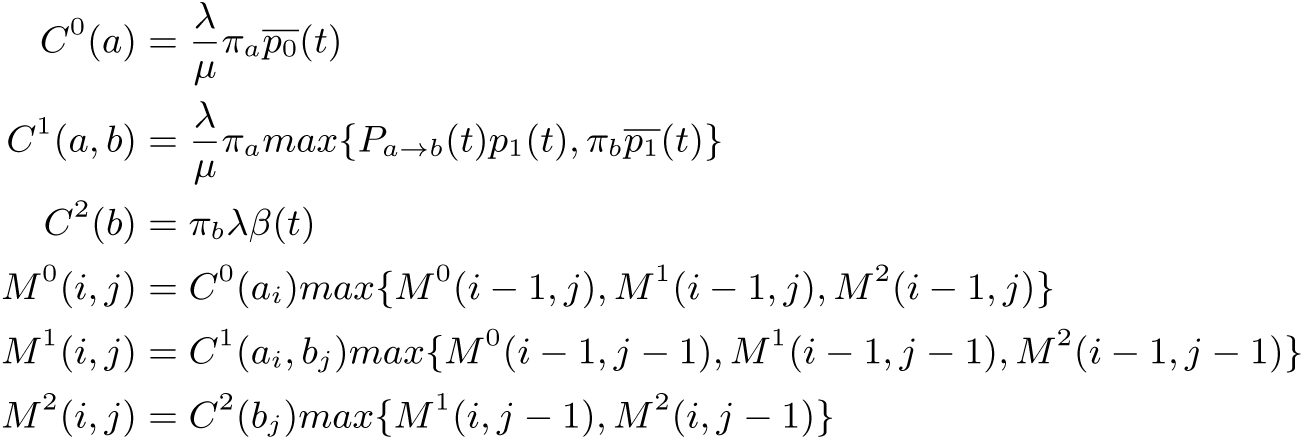
A re-formulation of the DP step using the memoized sub-problems (lookup tables) *C^i^.*

At a later point of the project, it became evident that the indexed loads from the memoized penalty matrices *C*^0^, *C*^1^, *C*^2^, as shown in Figure 5 caused a performance degradation. To address this problem, we tried to pre-compute a larger lookup table of vector-sized elements, that we indexed by a specific nucleotide permutation. Consider the following example for AVX vector intrinsics. We need to calculate a bijective mapping for a set of four nucleotides to an integer representing one of the 4^4^ = 256 possible permutations (e.g., a perfect hash) for *C*^0^ and *C*^2^, and one of the possible 4^2·4^ = 65536 permutations for *C*^1^ respectively. The mapping is implemented as base conversion from the set of 4 nucleotides to an unsigned integer via appropriate bit operations.

While this approach has exponential space requirements as a function of the vector width, using a lookup table of 65536 · 32 bytes = 2 MB for *C*^1^ for the AVX version of our code was still feasible. However, the performance evaluation revealed that too much time is spent to populate the table. Thus, we abandoned this approach.

### 3.4 Vectorization

We implemented a vectorized version for computing *M*^0^, *M*^1^, and *M*^2^ using `add`, `max`, `load`, and `store` operations for both SSE3 and AVX instructions. With these operations, we can process *n* matrix elements in parallel. Depending on the location of the vector on the anti-diagonal that is being processed, there will be 0 < *n* ≤ *V* valid elements to operate on per vector (with vector size *V* := 4 for AVX and *V* := 2 for SSE3).

Usually, data that has to be loaded into vectors needs to be aligned, that is, the starting address of the vector data needs to be a multiple of 16 or 32 bytes. This memory alignment allows to use the aligned versions of the vector `load` and `store` operations, which are faster than the respective unaligned operations. Although we ensured memory-aligned accesses to the majority of the data (as will be explained in Section 3.5), we consistently used unaligned vector `load` and `store` operations (`loadu_pd` and `storeu_pd`) in the SSE3 and AVX versions of our code. We did not observe a significant performance degradation when applying unaligned load intrinsics on aligned data.

To prevent our `store` operations from overwriting data at the boundaries of the matrices, we ensured that for vectors with size *n < V* (i.e., not completely filled/padded vectors) only *n* elements are written back into the matrices. For the SSE3 version, this can be achieved by only writing the lower part of the vector if its size is 1. For AVX, however, we covered all cases where *n < V* holds using the `maskstore_pd` intrinsic and an appropriate mask for all possibles values of *n* (i.e., 3, 2, and 1).

### 3.5 Data Alignment

If we want to vectorize along the anti-diagonals, accessing the memory in the DPMs when they are stored in row-, or column-major order decreases efficiency. This is because such a DP storage scheme does not allow to deploy efficient vector operations for loading the values from the matrices into vector registers. For such a matrix layout, the performance of the inner DP loop becomes heavily memory-bound. Therefore, we used the aforementioned layout by consecutive anti-diagonals together with the struct of arrays approach (see section 3.2), such that unaligned `load` and `store` operations for moving data directly to/from the anti-diagonal into vector registers can be utilized.

**Fig. 6.**
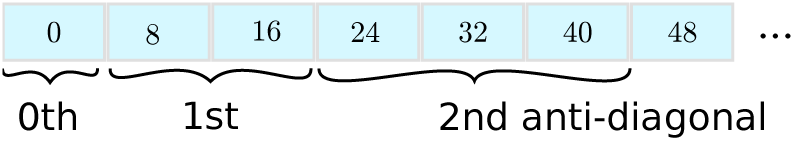
Shift of the byte offset of the anti-diagonals

Figure 6 shows why a 16/32 byte alignment for the starting addresses of the anti-diagonals can not be achieved when just storing anti-diagonals linearly. To solve this problem, we used padding, that is, we allocate some dummy entries such that each anti-diagonal starts at a 16/32 byte-aligned address. As a consequence we also had to modify the indexing function from Section 3.2 accordingly. The main idea is to store each anti-diagonal starting at a 16/32 byte-aligned address.

We implemented a scheme where the row with index 1 is memory-aligned, because the row with index 0 is initialized *prior* to entering the DP loop. As a consequence, each anti-diagonal is aligned to an “odd” address (*a −* 8, where *a* is the desired byte alignment, for instance, 32 − 8 = 24 bytes for AVX intrinsics).

In Figure 7, we show a snapshot of the required values for computing the antidiagonal elements with AVX intrinsics (vector length: 4 double precision values). In the current step we want to calculate the red elements. For calculating a single element, we require three elements from the two previous anti-diagonals. We need to access the element directly above (yellow) and left (blue) to the current element as well as the diagonal element above and to the left (green). The Figure illustrates that —omitting the asymptotically irrelevant boundary cases— one of the following four conditions holds:

1. red and blue start on an aligned address, yellow and green do not
2. red and yellow start on an aligned address, blue and green do not
3. yellow and green start on an aligned address, red and blue do not
4. red and green start on an aligned address, yellow and blue do not

**Fig. 7.**
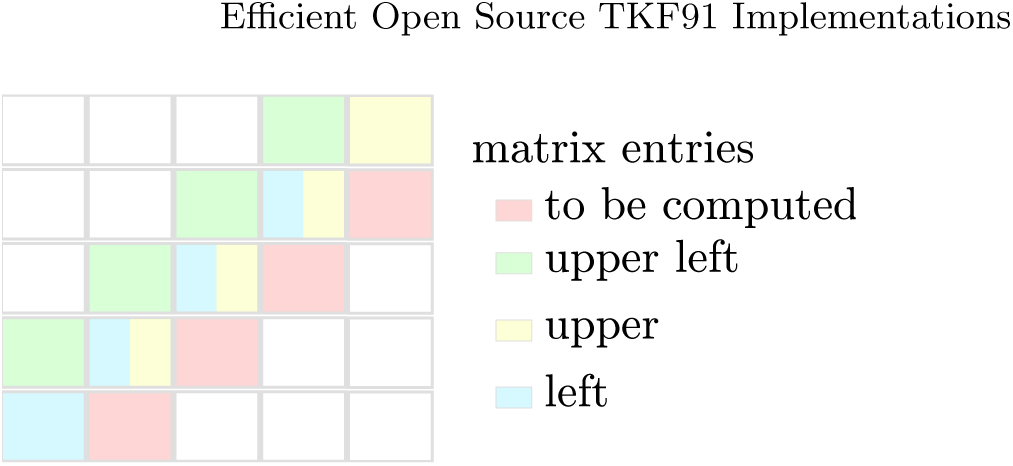
Outline of the data accesses required for calculating an anti-diagonal.

For the computation, we need to perform three `load` operations in total for the blue, yellow, and green elements as well as one `store` operation for the red elements. Since, based on empirical observations, *unaligned* `store` operations require more time than *unaligned* `load` operations, we ensured that either condition 1 or 2 above is always met. Thus, we implemented our code such that, the first condition is met for the opening and intermediate part and the second condition holds true for the closing part of the DP matrices.

## 4 Implementation of Team II

In the following, we first describe how we transformed the algorithm into logspace to prevent numerical underflow. This also allowed us to simplify the formulas. Thereafter, we report how we improved data locality by storing the matrix entries in a dedicated data structure. While we experimented with different vectorization techniques, it turned out that the fastest code did not rely on vectorization. We conclude, that we managed to simplify the sequential code to a point, where the vectorization overhead (see Section 4.3) exceeds the performance gains that can be achieved.

### 4.1 Mathematical Optimization - Basic Version

**Numerical Underflow Prevention** As the TKF91 algorithm performs successive multiplications of floating point numbers, preventing numerical underflow *is* a major issue. Underflow can occur, even for short input sequences with less than 100 nucleotides each. To address this issue we transformed all computations into log-space, to add logarithms of probabilities instead of multiplying probabilities. While it is still possible to experience numerical underflow, even after this transformation, we expect that this is highly unlikely to occur for practical (empirical) input data. We did not observe any numerical underflow for the range of input parameter values and sequence lengths (up to and including 10, 000 nucleotides) we tested.

**Simplifying the Formulas** We were able to omit redundant computations by re-using already computed values from previous matrix entries. For example, we observed that

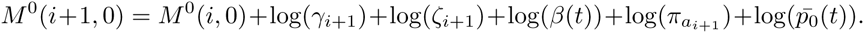

After replacing 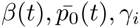, and *γ_i_* by *ζ_i_* their respective formulas, we noted that some terms appear multiple times. Operating in log-space allowed us to further simplify the formulas. Especially the logarithmic rules log(*a*b*) = log(*a*) + log(*b*) and log(*a/b*) = log(*a*) − log(*b*) allowed us to replace multiplications and divisions with additions and subtractions.

In the following formulas, 1 ≤ *i ≤ n* and 1 ≤ *j* ≤ *m*, if not stated otherwise.

*Matrix Initialization*

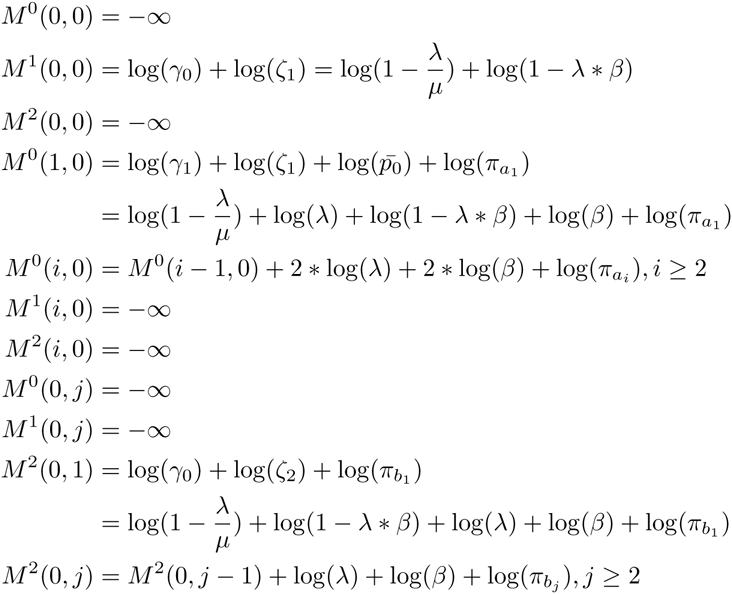

*Further Initialization*

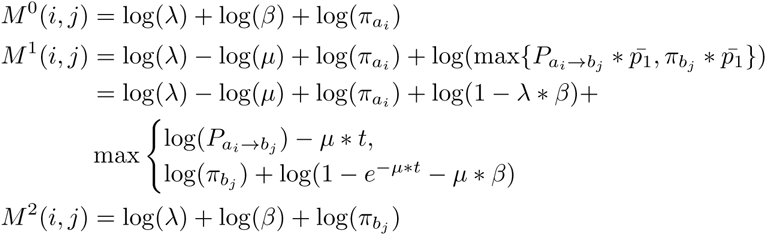

*Dynamic Programming Step*

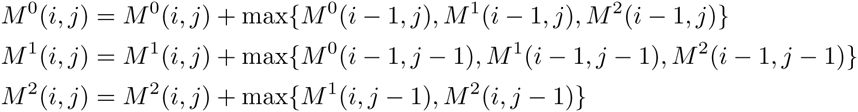

**Pre-computing the Logarithms** In general, replacing a multiplication by two logarithms and an addition is more costly. However, we managed to circumvent the additional computational cost of log space. This is because our formulas above contain only 25 different logarithm invocations that only depend on given, constant input parameters. Hence, all these logarithms can be pre-computed.

A major disadvantage of logarithmic transformations is that errors in the logarithm computation accumulate over a sequence of operations. To further investigate this, we used the high precision mathematical library of the `boost`-framework [1]. It offers datatypes that dynamically adapt their floating point precision to avoid numerical errors as well as under-/overflow. Our initial idea was to use this library only for pre-computing the logarithms, and thereby reduce the induced runtime overhead. However, because we converted these arbitrary precision datatypes back into standard double precision floating point values (using `convert_to<double=()`) for increasing the speed of the actual DP calculations, we did not observe increased precision. Therefore, we abandoned this path.

In the rest of the manuscript we refer to the code obtained by applying the above techniques and transformations as the basic version.

### 4.2 Matrix Storage Schemes - Improved Version

**Matrix as Array** In the first, naive implementation, we stored each matrix (*M*^0^, *M*^1^, *M*^2^) in a separate array, using row-major order. The matrix entry at position (*i, j*) is stored at index position *i* * (*m* + 1) + *j* in the array (see Figure 8).

To illustrate the shortcomings of this approach, consider the following example: assume that six matrix rows fit into one cache line. Then, performing one iteration of the inner DP loop requires loading three cache lines, one per DPM. This is depicted in part a) of Figure 9. If we further assume a cache capacity of only two cache lines, every iteration would then force one of the lines to be swapped out and decrease memory efficiency.

**Fig. 8.**
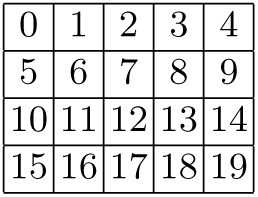
Row-major indexing

**Fig. 9.**
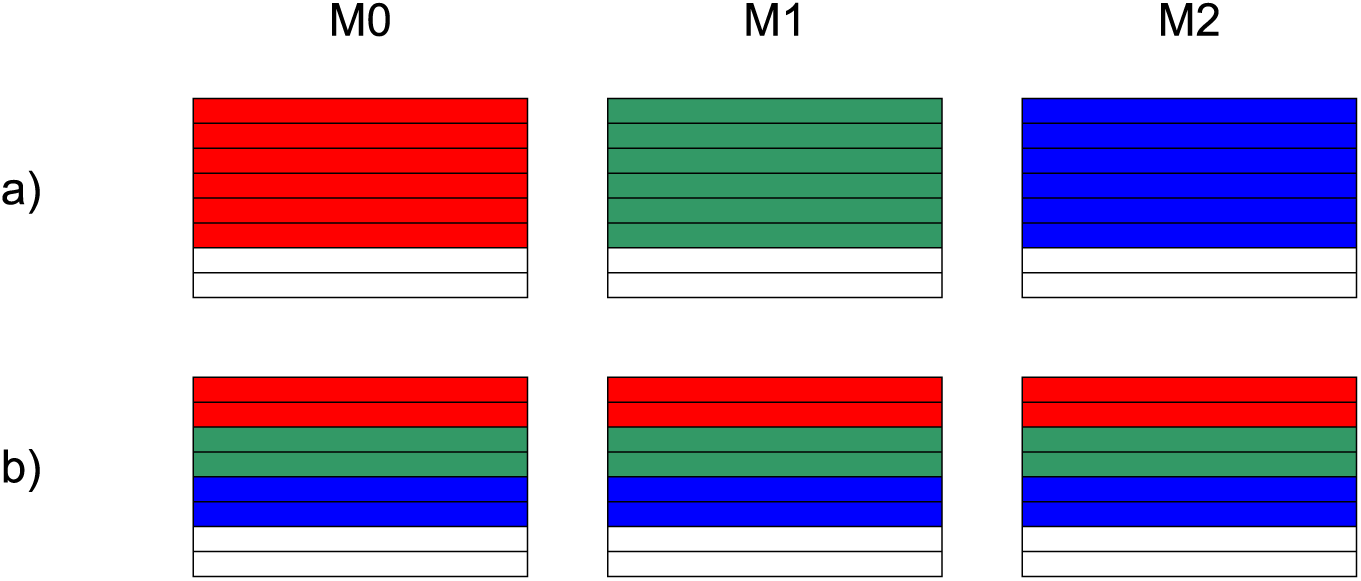
Relation of cache lines and matrices, using the method of allocating each matrix separately(**a**) and the Array-of-Structs method (**b**). Each color represents a distinct cache line.

**Array-of-Structs Data Structure** The dynamic programming step (see Algorithm 1) accesses the matrices *M*^0^, *M*^1^ and *M*^2^ at the same index position to determine the maximum entry.

Thus, we improved data locality by storing the entries from the three matrices located at identical index positions contiguously in memory. For this, we implemented a data structure called `MatrixEntry` that consists of three double values: *m*_0_, *m*_1_ and *m*_2_. Then, we store the matrices linearly (see Section 4.2). In contrast to the initial approach, we only store one array and each array element now is a `MatrixEntry` struct instead of a simple `double` value (see Figure 10 and *SoA* as defined by team I in Section 3.2).

Let us now consider the example in Figure 9 again. With six matrix lines filling one cache line, a cache line now contains data from all three matrices. Consequently, all operations for a single DP cell update will access at most two cache lines. If we assume a cache capacity of two cache lines again, cache misses will now only occur when traversing row boundaries.

By using the `perf stat` tool, we found that using the `MatrixEntry` data structure indeed reduced the number of page faults which we use as a proxy for cache efficiency. The number of page faults was a sufficient proxy for predicting cache-related performance. Thus, we did not deploy more elaborate cache simulators such as `cachegrind`.

#### Algorithm 1

The dynamic programming step, row-major version, where *CO*(*i, j*) maps the DP coordinates *i* and *j* to the linear index in the array of structs data structure.

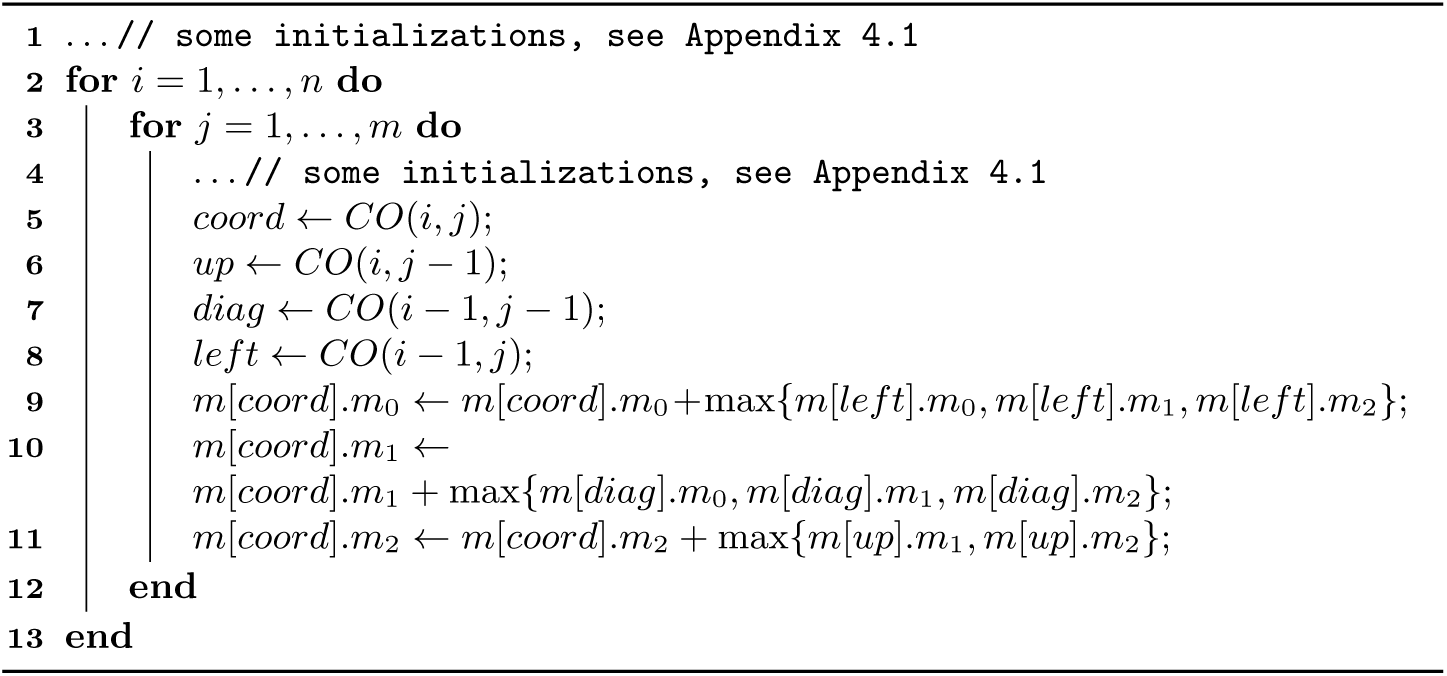

**Fig. 10.**
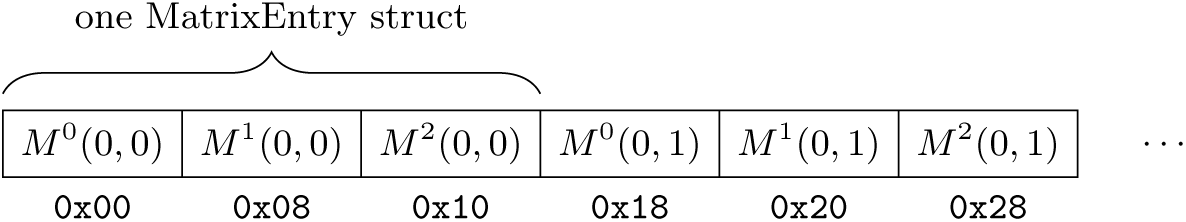
Array-of-Structs data structure for storing the three matrices in memory. Each `MatrixEntry` element stores the entries of a single index for all three matrices. The structs are stored contiguously in row-major order.

**Alternative Storage Schemes** Since each DP cell update needs to access the top, left, and upper-diagonal elements of the matrices, storing the matrix anti-diagonals linearly (see Figure 11 and considerations by Team I) will minimize cache misses. Finding an inexpensive-to-compute closed formula that maps the index position (*i, j*) representing the row and column of a matrix to wave-front-coordinates turned out to be challenging (see also discussion by Team I in Section 3.2). The index of the diagonal is *i + j*, the sum of the row index and the column index. Determining the number of elements on the anti-diagonal and especially the correct position along the anti-diagonal proved more difficult though. The indexing formulas we tested were too computationally expensive and required more computations than the actual cell updates. As an alternative, we tried storing pre-computed index mappings in an additional matrix. However, this slowed down the program as computing the mapped indices required more arithmetic operations than the actual TKF91 algorithm. Figure 12 illustrates the main problem which has to be solved for efficiently indexing-diagonals: Given an wave-front index *k*, how do we obtain the indices for the upper, upper-left-diagonal, and left element of the matrix? While the required offsets are constant for a single anti-diagonal, they change for successive anti-diagonals.

**Fig. 11.**
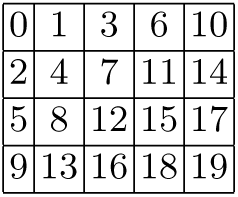
Wave-front indexing

**Fig. 12.**
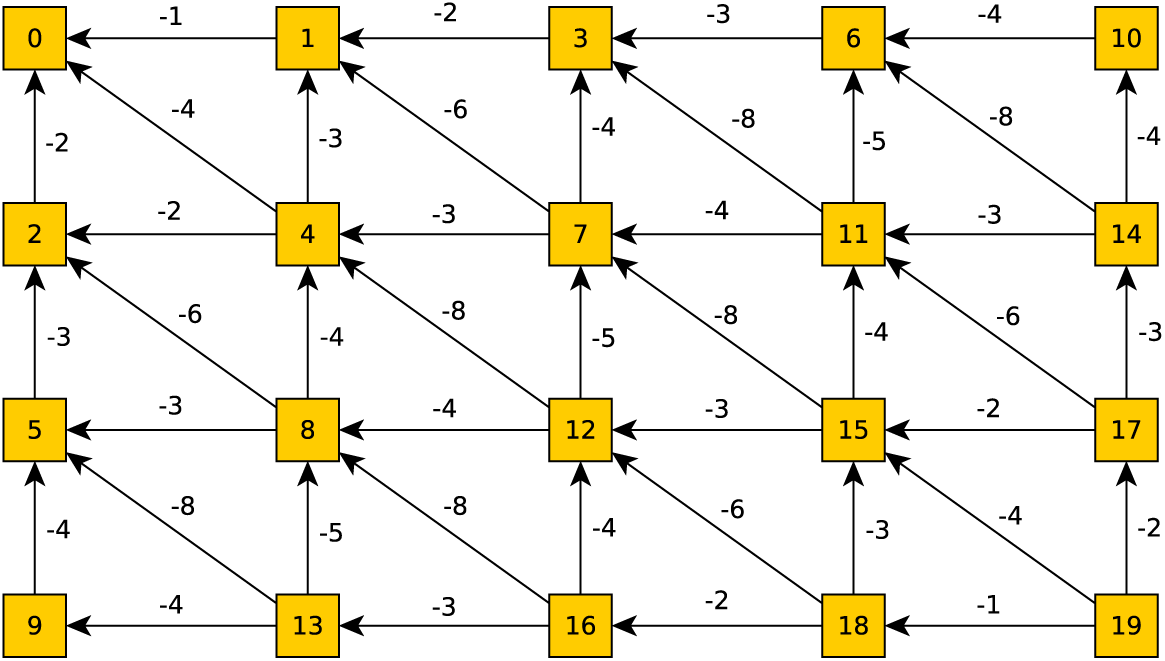
Offsets in wave-front/anti-diagonal indexing

We denote the code obtained by applying the above storage scheme, the improved version.

### 4.3 Vectorization Attempts

During code development we conducted a partial vectorization for two of our implementations: the basic log-space implementation (Section 4.1) and the improved log-space implementation, using a more cache-efficient data structure to store the matrices (Section 4.2).

In both cases we mainly focused on vectorizing the code that initializes the three matrices. This is because our analyses with `Valgrind callgrind` revealed, that this part of the code required 75 − 80% of overall runtime. Additionally, as the initializations are completely independent of each other, this part should be straight-forward to vectorize.

Several code transformations were necessary to vectorize the code. First, we had to ensure that vector `load` and `store` operations are performed on correctly aligned memory addresses (see discussion by Team I). Secondly, we needed to devise a strategy for iterating over matrices and handling matrix sizes that are not multiples of the vector width. Finally, we needed to store calculation terms (e.g., log(*π*)’s or 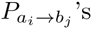), in appropriate vector types and had to replace all basic arithmetic operations by their vectorized counterparts.

**Log-space implementation** For the vectorization of our basic version (see Section 4.1), we implemented an appropriate memory alignment by using the `__attribute__(aligned(size))` attribute for stack allocations, and the respective `posix_memalign` function for heap allocations.

We also changed the iterations through the matrices to start at entry 0 of each row, as starting iterations at position 1 would result in memory accesses at incorrectly aligned addresses. Furthermore, as we store the matrix rows contiguously in memory, we had to pad the rows to ensure a correct alignment of the first entry in each row.

The strategy we chose to deal with row dimensions that are not multiples of the vector width is straight-forward. For each row, we compute as much as possible via vector intrinsics and the remainder sequentially.

**Version using Array of Structs data structure** While the strategy for dealing with row dimensions that do not fit vector widths remained the same in the improved version (introduced in Section 4.2) of our program, we had to apply several modifications to be able to use vector intrinsics in conjunction with this more cache-efficient data structure.

As Figure 10 shows, using the Array-of-Structs scheme results in a noncontiguous storage of row entries. As a consequence, loading and storing vector registers is not straight-forward. To this end, we implemented load and store operations that fit our data structure. These operations rely on vector functions that load and store single double precision floating point values. An advantage of this is that we do not need to enforce correct data alignment any more. However, a single `_mm_load_pd` on contiguous memory is substantially faster than a series of `_mm_loadl_pd` and `_mm_loadh_pd` operations on non-contiguous memory locations using SSE3 intrinsics.

**Thoughts on wave-front vectorization** Our two log-space versions of TKF91 reduce the DP cell update calculations that determine the largest of two or three values. In turn, this means that the relative amount of time spent for DP cell updates is almost negligible. Nevertheless, we invested time to explore potential vectorized wave-front versions of the program.

Initially, as Team I, we investigated appropriate DP cell storage and access schemes, to store the data contiguously with respect to the wave-front parallelization data access pattern. We found that, indexing a cell required 20–30 arithmetic operations as opposed to only 2–3 for the actual cell update. Thus, merely indexing a cell, using the standard row and column coordinates, requires an excessive amount of computations. Our preliminary benchmarks yielded absolutely no performance improvement for the vectorized versions of the basic as well as the improved versions of our code. Therefore, we abandoned the vectorized wave-front approach.

## 5 Evaluation and Testing

We initially describe the benchmark data we used (Section 5.1) as well as the experimental setup. Then we present the runtime results for teams I and II. We conclude with a thorough analysis of the impact of the logarithm implementation used in Section 5.3.

### 5.1 Test Data and Experimental Setup

For testing and performance assessment, we used empirical benchmark datasets. We download multiple sequence alignments (MSAs) from http://goo.gl/nlD4nb that were used in [2]. From each of the six MSAs, we selected ten sequences (e.g., for an MSA with 60 sequences we extracted the sequences with indices 0, 6, 12,…, 54), and initially dis-aligned the sequences (removed all MSA gaps). Then, we computed all 45 possible pairwise TKF91 alignments between these ten sequences. As input parameter values, we used λ := 1.0, *μ* := 2.0, *τ* := 0.1, *π* := (0.27, 0.24, 0.26, 0.23). We empirically determined that using median of five samples, where each sample corresponds to the average runtime for ten TKF91 executions (i.e., a total of 5 * 10 * 45 = 2250 pair-wise TKF91 alignments per dataset) yielded stable runtime estimates.

As test platform we used the aforementioned reference hardware (see Section 2.1).

In Sections 5.2 and 5.3 the two student teams present results of specific performance and accuracy issues they investigated in detail. In Section 5.4 we compare the runtimes of the implementations developed by teams I and II.

**Table 1.** Team I benchmark results for the SoA, AoS, and AoSoA storage schemes. The fastest execution time is shown in bold font.

| ms per iteration<br>(compared to best) | SoA | AoS | AoSoA |
| --- | --- | --- | --- |
| BDNF | 83.1 (1.01) | **82.2 (1.00)** | 85.1 (1.04) |
| cytb | **262.9 (1.00)** | 290.7 (1.11) | 296.5 (1.13) |
| RAG1 | 209.1 (1.02) | **205.6 (1.00)** | 209.4 (1.02) |
| RAG2 | **246.1 (1.00)** | 250.9 (1.02) | 258.3 (1.05) |
| RBP3 | **346.9 (1.00)** | 363.5 (1.05) | 376.2 (1.08) |
| vWF | **439.6 (1.00)** | 441.0 (1.00) | 462.7 (1.05) |

### 5.2 Results of Team I

As mentioned in Section 3.5, we performed various performance tests using this benchmark suite. We tested different vector load/store schemes and measured the run times of the SoA, AoS, and AoSoA storage schemes. We show the results of the SoA versus AoS versus AoSoA runtime experiments in Table 5.1. As already mentioned in Section 3.5, the SoA approach performs best.

### 5.3 Results of Team II

We first assess the performance and impact on the output of distinct logarithm implementations (Section 5.3). Thereafter, we measure the runtime of our code in Section 5.3.

**Different Logarithm Libraries** In addition to the standard `log` function (from C++ `<math.h>`), we also used the `crlibm` library by Daramy *et al.* [3]. It includes versions of the logarithm function, which allow to explicitly specify the rounding strategy, (e.g., rounding up, rounding down, towards zero, or towards the nearest integer), to be used.

To assess the numerical stability of our alignments, we computed the edit distances between alignments calculated using distinct logarithm implementations to a reference alignment using the `Boost.Multiprecision` library. The edit distance quantifies the difference between two strings. It calculates the minimum cost sequence of string edit operations required (insertions, deletions, and substitutions; note that these are string edit operations not related with the TKF91 model), to transform one string into the other. We assigned a cost of 1 to insertions, deletions, and substitutions of single letters. For our tests, we aligned all possible pairs of sequences from the BDNF MSA in the aforementioned benchmark data (see Section 5.1).

For the alignments, we used λ := 1, *μ* := 2, *π* := (0.27, 0.24, 0.26, 0.23) and *τ* := 0.1.

In particular the `log_ru` function from the `crlibm` library, which explicitly rounds up, produces a different alignment than the `Boost.Multiprecision` version (see Table 2). However, the computed likelihood scores were highly similar and edit distance between the alignments was minimal between the `crlibm` and `Boost.Multiprecision` versions (see Figure 13). We finally decided to use the `log_ru` function from the crlibm library because, in most cases, it yielded the same results as the sample solution provided by our instructors (http://www.exelixis-lab.org/web/teaching/practical15/scaledCode/tkf91_scaling.tar.gz).

**Fig. 13.**
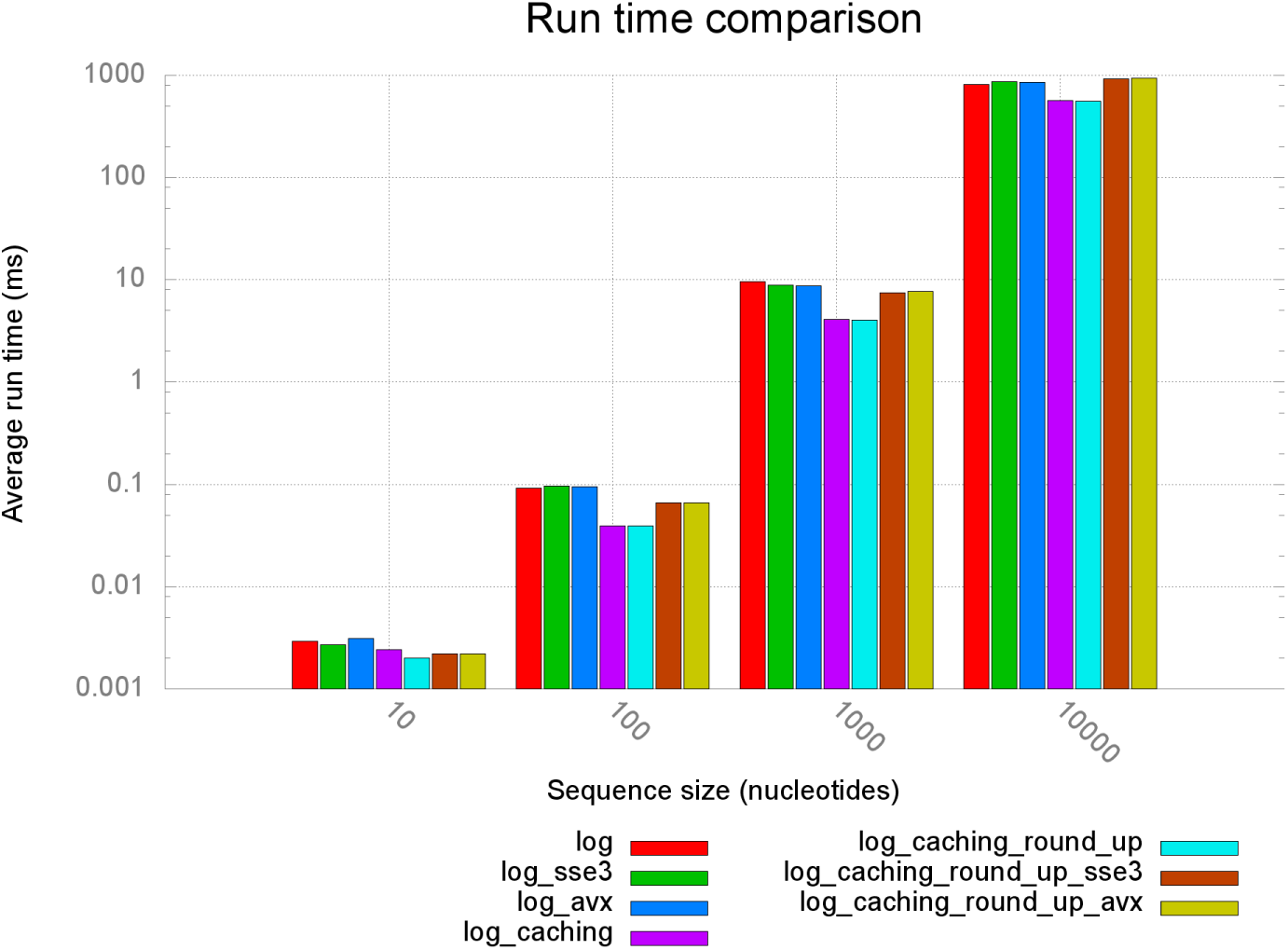
Team II: Numerical alignment differences for input parameters *π* : = (0.25, 0.25, 0.25, 0.25), λ := 1, *μ* := 2, *τ* := 0.1

**Table 2.**
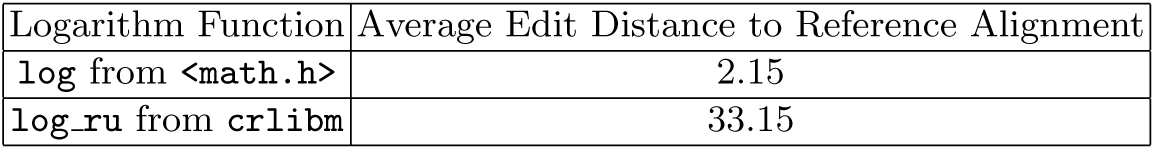
Team II: average edit distances from the `Boost.Multiprecision` reference alignment for different logarithm implementations.

| Logarithm Function | Average Edit Distance to Reference Alignment |
| --- | --- |
| `log` from `<math.h>` | 2.15 |
| `log_ru` from `crlibm` | 33.15 |

**Runtime Measurements** For measuring runtimes during code development, we used four pairs of randomly generated sequences consisting of 10, 100, 1000, and 10000 nucleotides, respectively.

We measured the execution times around the kernel of the program, that is, excluding any I/O required to load parameters and input sequences, or the output of the program. The allocation and initialization of the three DPMs form part of the kernel and were therefore included in our measurements. We executed the kernel multiple times for each dataset and subsequently averaged the runtimes. To balance accuracy and overall benchmark time (see Table 3) we used distinct numbers of kernel invocations, depending on the dataset size.

The results of these runs are summarized in Figure 14 for 7 distinct version of our code using different logarithm implementations, SSE3 and AVX intrinsics for initializing the DPMs, and using the more cache-efficient array of structs storage scheme (denoted by `_caching`).

**Table 3.** Team II: number of runs per sequence length

| Length of Sequences | Number of Runs |
| --- | --- |
| 10 | 10000 |
| 100 | 1000 |
| 1000 | 100 |
| 10000 | 10 |

**Fig. 14.**
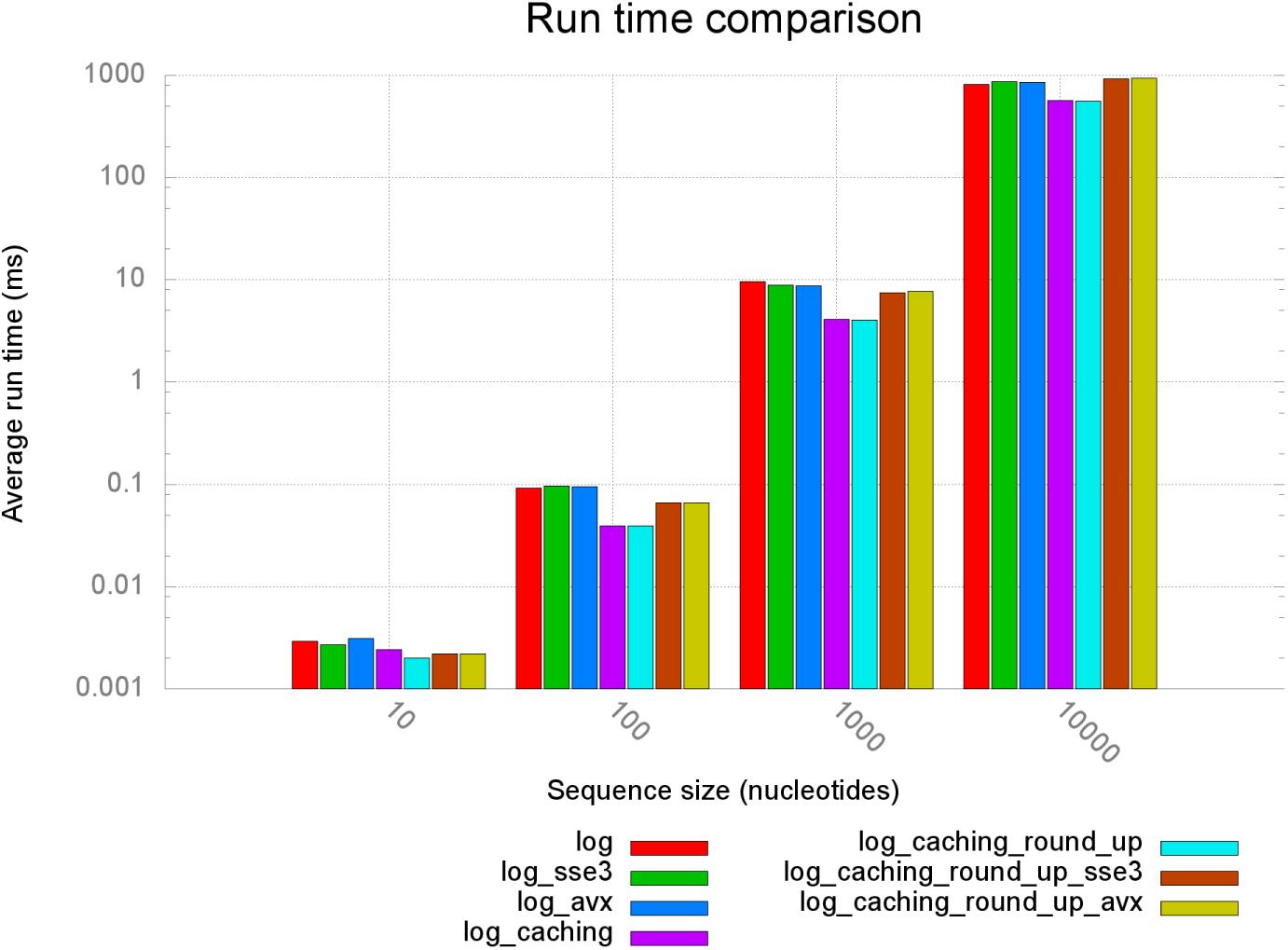
Team II: comparison of average runtimes of different implementations. `sse3` and `avx` post-fixes denote the vectorized versions of the programs (see Section 4.3). With `log` we denote the log-space transformed versions of TKF91 (Section 4.1). Versions using an arrays of structs to store matrices (Section 4.2) are denoted by `caching`; `round_up` denotes versions relying on the `crlibm` library logarithm function `log_ru` (Section 5.3).

### 5.4 Performance: Team I versus Team II

To test performance we deployed the Celero benchmark suite that can directly measure the run-time (see https://github.com/DigitalInBlue/Celero) of a single function call. We used the Celero suite to compare the performance of the codes developed by teams I and II on the aforementioned benchmark data set (see Section 5.1). In Table 5.4 we depict the runtimes for a single kernel invocation in milliseconds as well as the runtime ratio with respect to the fastest implementation. Note that, the run times correspond to the accumulated execution time over all 45 possible pair-wise alignments between the 10 sequences we selected from each dataset. The performance comparison was set up jointly by the two teams to guarantee a fair comparison.

**Table 4.** Performance comparison of TKF91 implementations by teams I & II.

| ms per iteration<br>(compared to best) | Team I | Team II |
| --- | --- | --- |
| BDNF | 83.1 (1.50) | **55.4 (1.00)** |
| cytb | 262.9 (1.25) | **209.9 (1.00)** |
| RAG1 | 209.1 (1.42) | **147.3 (1.00)** |
| RAG2 | 246.1 (1.28) | **192.0 (1.00)** |
| RBP3 | 346.9 (1.25) | **277.6 (1.00)** |
| vWF | 439.6 (1.25) | **352.2 (1.00)** |

## 6 Teaching Results

### 6.1 What did Team I learn?

While the assigned task was straightforward and very manageable in principle, the implementation details were tricky. Preventing floating point underflow while, at the same time, optimizing performance was challenging. This is because, such technical problems are frequently ignored in other programming practicals at KIT that focus on functionality. Furthermore, vectorizing with SIMD intrinsics required some rethinking and re-engineering because we needed to identify a suitable memory layout, optimize data alignment, and deal with boundary conditions (padding). While it was satisfying to address and solve problems progressively, there were always additional ideas to further improve the code (w.r.t. performance and design), which made prioritizing tasks important. We enjoyed the satisfaction of gaining yet another percent of execution time in combination with working with SIMD instructions that were not familiar to us.

The basic problem was clearly outlined. There was enough freedom and time left to work on improving our solution. The scheduled project milestones turned procrastination and last minute work into a non-issue. Most surprisingly, we were always on schedule.

### 6.2 What did Team II learn?

For this assignment we had to tackle two major problems: preventing numerical underflow and pursuing the elusive “most efficient” implementation. In the early stages, we came up with the idea of transforming the algorithm into log-space to solve the numerical issues. This did not only prove to represent an efficient solution, but also allowed us to further simplify the formulas and to pre-compute the vast majority of terms. In this context we also discovered deviations in the output alignments that depend on the specific implementation of the logarithm function being used. We therefore used the `crlibm` library of mathematical functions, that allowed us to assess the impact of logarithm functions with distinct rounding strategies on the final result.

To optimize the code we also used SSE3 and AVX intrinsics to vectorize the most work-intensive portions of the code. While we were unable to produce a faster vectorized code, we did learn how to use vector intrinsics and how to deal with the associated pitfalls. The largest performance gain was attained via a more cache-efficient data structure. For this, we used profiling tools such as `perf` and `clang` for the first time. Finally, we concluded that `C++` is appropriate for HPC projects, provided that, the problem and data-structures at hand are well understood.

## 7 Conclusion

We have implemented and made available two independent, highly optimized open-source implementations of the TKF91 algorithm for statistical pair-wise sequence alignment. We thoroughly assessed their performance and investigated aspects such as differing alignment results because of slight numerical deviations in logarithm implementations. The implementations by teams I and II are substantially different in the approach they take for optimizing program performance on standard x86 architectures. We proposed several methods for storing and addressing the three dynamic programming matrices. In addition, we show how the original equations can be simplified to (i) avoid numerical underflow issues and (ii) save a substantial amount of computations.

We make the source code available in the hope that, it will be useful to the community. In particular, some of the wave-front vectorization approaches might be applied to the more complex statistical alignment kernels used in programs such as BaliPhy [7].

Finally, the task and code at hand can be used to design similar bioinformatics programming practicals. For instance, one might ask students to try and come up with a faster implementation or use the existing implementations as library functions in some larger and more complex programming project.

## Footnotes

3 Wall -Wextra -Wredundant-decls -Wswitch-default -Wimport –Wno –int –to -pointer -cast -Wbad-function-cast -Wmissing-declarations -Wmissing-prototypes -Wnested -externs -Wstrict-prototypes -Wformat-nonliteral –Wundef

4 Weverything pedantic

